# Altered functional interactions between neurons in primary visual cortex of macaque monkeys with experimental amblyopia

**DOI:** 10.1101/604009

**Authors:** Katerina Acar, Lynne Kiorpes, J. Anthony Movshon, Matthew A. Smith

## Abstract

Amblyopia, a disorder in which vision through one of the eyes is degraded, arises because of defective processing of information by the visual system. Amblyopia often develops in humans after early misalignment of the eyes (strabismus), and can be simulated in macaque monkeys by artificially inducing strabismus. In such amblyopic animals, single-unit responses in primary visual cortex (V1) are appreciably reduced when evoked by the amblyopic eye compared to the other (fellow) eye. However, this degradation in single V1 neuron responsivity is not commensurate with the marked losses in visual sensitivity and resolution measured behaviorally. Here we explored the idea that changes in patterns of coordinated activity across populations of V1 neurons may contribute to degraded visual representations in amblyopia, potentially making it more difficult to read out evoked activity to support perceptual decisions. We studied the visually-evoked activity of V1 neuronal populations in three macaques (*M. nemestrina*) with strabismic amblyopia and in one control. Activity driven through the amblyopic eye was diminished, and these responses also showed more interneuronal correlation at all stimulus contrasts than responses driven through the fellow eye or responses in the control. A decoding analysis showed that responses driven through the amblyopic eye carried less visual information than other responses. Our results suggest that part of the reduced visual capacity of amblyopes may be due to changes in the patterns of functional interaction among neurons in V1.

**New and noteworthy:** Amblyopia is a developmental disorder of visual processing that reduces visual function and changes the visual responses of cortical neurons in macaque monkeys. The neuronal and behavioral changes are not always well correlated. We found that the interactions among neurons in the visual cortex of monkeys with amblyopia are also altered. These changes may contribute to amblyopic visual deficits by diminishing the amount of information relayed by neuronal populations driven by the amblyopic eye.

## Introduction

Normal visual system development is dependent on having unobstructed and balanced binocular visual experience during early life. Amblyopia is a disorder of the visual system which often arises when visual input through the two eyes is imbalanced, most commonly through a misalignment of the two eyes (strabismus) or anisometropia (unilateral blur), during a critical window for development. Amblyopic individuals show major impairments in basic spatial vision in the affected eye, including decreased visual acuity and diminished contrast sensitivity that is often particularly acute at high spatial frequencies (*Baker et al. 2008; Bradley and Freeman 1981; Hess and Howell 1977; Levi and Harwerth 1977; McKee et al. 2003; for reviews see Asper et al. 2000; Levi 2013; Meier and Giaschi 2017*). Furthermore, several studies suggest that amblyopia is detrimental to cognitive processes that rely on higher visual system function, namely contour integration, global motion sensitivity, visual decision-making, and visual attention (*Farzin and Norcia 2011; Hou et al. 2016; Kozma and Kiorpes 2003; Kiorpes et al. 2006; Levi et al. 2007; Meier et al. 2016; Pham et al. 2018; Rislove et al. 2010; for reviews see Hamm et al. 2014; Kiorpes 2006, 2016; Meier and Giaschi 2017*). Deficits in amblyopic vision originate from altered neural activity in the primary visual cortex (V1), and cortical areas downstream of V1, rather than from abnormalities in the eye or the visual thalamus (*Bi et al. 2011; Blakemore and Vital-Durand 1986; Kiorpes et al. 1998; Movshon et al. 1987; Shooner et al. 2015; Smith et al. 1997; Tao et al. 2014; Wiesel 1982; for reviews see Kiorpes 2016; Kiorpes and Daw 2018; Levi 2013*).

Previous studies using macaque models of amblyopia provide evidence for some functional reorganization of ocular dominance in amblyopic V1 (*Adams et al. 2015; Crawford et al. 1989; Crawford and Harwerth 2004; Fenstemaker et al. 2001; Hendrickson et al. 1987; Horton et al. 1997; LeVay et al. 1980; Tychsen et al. 1992, 1997, 2004*), including a significant loss in the proportion of binocularly activated cells and – in severe amblyopia – a reduced proportion of neurons that respond to amblyopic eye stimulation (*Kiorpes et al. 1998; (in cat) Schröder et al. 2002; Shooner et al. 2015; Smith et al. 1997)*. Additionally, several studies report changes in spatial frequency tuning, as well as a loss of contrast sensitivity in some V1 neurons that receive input from the amblyopic eye in monkeys (*Kiorpes et al. 1998; Movshon et al. 1987*) and in cats (*Chino et al. 1983; Crewther and Crewther 1990).* Overall, these changes in the functional properties of V1 neurons suggest that the representation of visual input from the amblyopic eye across the cortical neuronal population is distorted.

Early studies on the neural basis of amblyopia hypothesized that the perceptual deficits in amblyopes arise directly from corresponding losses in responsivity of single neurons in primary visual cortex (*Wiesel and Hubel 1963; Wiesel 1982*). However, it is now clear that the magnitude of these single neuron changes cannot account for the entirety of spatial vision deficits revealed by behavioral assessments of amblyopes (*Bi et al. 2011; Kiorpes et al. 1998; Shooner et al. 2015*). There are two additional neurophysiological mechanisms that could contribute to amblyopia: (1) neural deficits more profound than those seen in V1 may arise in downstream visual areas (*Bi et al. 2011; El-Shamayleh et al. 2010*; Kiorpes et al. 1998, 2016; Tao et al. 2014; *Wang et al*. 2017) and (2) impaired visual representation might result from changes in the structure of activity in populations of V1 neurons (*Kiorpes 2016; Roelfsema et al. 1994; Shooner et al. 2015*).

Here we seek evidence for this second mechanism, and investigate whether activity correlations between neurons are altered in amblyopic V1 during visual stimulus processing. We recorded from populations of V1 neurons in macaque monkeys that had developed amblyopia as a result of surgically-induced strabismus (*as in Kiorpes et al. 1998*). We measured correlation in the trial-to-trial variability (hereafter referred to as “correlation”) in the responses of pairs of neurons to an identical visual stimulus presented to either the non-amblyopic (fellow) or amblyopic, deviating eye. Similar to the firing rate of single neurons, the strength of correlated variability in normal visual cortex has been shown to change due to a number of factors, including the contrast of a visual stimulus (*Kohn and Smith 2005*), the animal’s attentional state (*Cohen and Maunsell 2009; Mitchell et al. 2009; Snyder et al. 2016*), and over the course of perceptual learning (*Gu et al. 2011; Ni et al. 2018).* In our experiments, comparing correlation measurements for stimuli presented to the two eyes allowed us to determine whether the functional circuitry used for processing amblyopic eye visual input is altered compared to that supporting fellow eye processing. We found that correlation indeed changes depending on which eye receives the visual stimulus, an effect that was not present in a control animal. Overall, stimuli presented to the amblyopic eye evoked correlations that were more prominent in pairs of neurons with similar orientation tuning and eye preference. When stimulus contrast was increased, pairs of neurons driven through the fellow eye tended to decorrelate, whereas the high levels of correlation remained elevated for neurons driven by the affected eye. Our findings are consistent with the hypothesis that the abnormalities in amblyopic vision may in part be explained by changes in the strength and pattern of functional interactions among neurons in primary visual cortex.

## Materials and Methods

### Subjects

We studied four adult macaque monkeys (*Macaca nemestrina*), three female and one male. One animal remained a visually normal, untreated control while three of the animals developed strabismic amblyopia as a result of surgical intervention at 2-3 weeks of age. Specifically we resected the medial rectus muscle and transected the lateral rectus muscle of one eye in order to induce strabismus. All of the animals underwent behavioral testing to verify the presence or absence of amblyopia. All procedures were approved by the Institutional Animal Care and Use Committee of New York University and were in compliance with the guidelines set forth in the United States Public Health Service Guide for the Care and Use of Laboratory Animals.

### Behavioral testing

We tested the visual sensitivity of each animal by evaluating their performance on a spatial two-alternative forced-choice detection task. Behavioral testing was conducted at the age of 1.5 years or older, and the acute experiments took place at the age of 7 years or older. On each trial in this task, a sinusoidal grating was presented on the left or the right side of a computer screen while the animal freely viewed the screen. The animal had to correctly indicate the location of the grating stimulus by pressing the corresponding lever in order to receive a juice reward. The gratings varied in spatial frequency and contrast level: we tested 5 contrast levels at each of 3-6 different spatial frequencies and collected at least 40 repeats of each stimulus combination. We then determined the lowest contrast the animal could detect at each spatial frequency (threshold contrast) and constructed contrast sensitivity functions for each animal’s right and left eyes. A detailed account of the procedures we used for behavioral assessment in this study can be found in previous reports (*Kiorpes et al. 1999; Kozma and Kiorpes 2003*).

### Electrophysiological recording

The techniques we used for acute physiological recordings have been described in detail previously (*Smith and Kohn 2008*). Briefly, anesthesia was induced with ketamine HCl (10 mg/kg) and animals were maintained during preparatory surgery with isoflurane (1.5-2.5% in 95% O2). Anesthesia during recordings was maintained with continuous administration of sufentanil citrate (6-18 μg/kg/hr, adjusted as needed for each animal). Vecuronium bromide (Norcuron, 0.1 mg/kg/hr) was used to suppress eye movements and ensure stable eye position during visual stimulation and recordings. Drugs were administrated in normosol with dextrose (2.5%) to maintain physiological ion balance. Physiological signs (ECG, blood pressure, SpO2, end-tidal CO2, EEG, temperature, and urinary output and osmolarity) were continuously monitored to ensure adequate anesthesia and animal well-being. Temperature was maintained at 36-37 C°.

Recordings of neural activity were made from 100-electrode “Utah” arrays (Blackrock Microsystems) using methods reported previously (*Kelly et al. 2007; Smith and Kohn 2008*). Each array was composed of a 10×10 grid of 1 mm long silicon microelectrodes, spaced by 400 um (16 mm^2^ recording area). Each microelectrode in the array typically had an impedance of 200-800 kOhm (measured with a 1 kHz sinusoidal current), and signals were amplified and bandpass filtered (250 Hz to 7.5 kHz) by a Blackrock Microsystems Cerebus system. We targeted the superficial layers by inserting the arrays 0.6 mm into cortex using a pneumatic insertion device (*Rousche and Normann 1992*).

Our full data set consisted of acute recordings from 7 microelectrode arrays across 3 amblyopic macaque monkeys and 4 arrays in 1 control monkey. One of the amblyopic animals (EM 640) had 4 array implants (3, 8, 14 and 51 neurons); one (JS 579) had 2 array implants (34 and 68 neurons), and the third (HN 580) had 1 array implant (30 neurons). The control animal had 4 implants (4, 7, 6, and 16 neurons). For animals with multiple implants in a single hemisphere, the array was removed and shifted to a different, non-overlapping region of cortex prior to reimplantation. We did not notice any substantial differences in recording quality across arrays moved to different locations. Arrays were inserted within a 10 mm craniotomy made in the skull, centered 10 mm lateral to the midline and 10 mm posterior to the lunate sulcus. The resulting receptive fields lay within 5° of the fovea.

### Visual stimulation

We presented stimuli on a gamma-corrected CRT monitor (Eizo T966), with spatial resolution 1280 × 960 pixels, temporal resolution 120 Hz, and mean luminance 40 cd/m^2^. Viewing distance was 1.14 m or 2.28 m. Stimuli were generated using an Apple Macintosh computer running Expo (http://corevision.cns.nyu.edu).

We used a binocular mirror system to align each eye’s fovea on separate locations on the display monitor, so that stimuli presented in the field of view of one eye did not encroach on the field of view of the other eye. This setup enabled us to show stimuli to the receptive fields for the right and left eye independently. We mapped the neurons’ spatial receptive fields by presenting small, drifting gratings (0.6 degrees; 250 ms duration) at a range of spatial positions in order to ensure accurate placement of visual stimuli within the recorded neurons’ receptive fields. During experimental sessions, we presented full-contrast drifting sinusoidal gratings at 12 orientations spaced equally (30°) in the field of view of either the right or the left eye on alternating trials. Each stimulus was 8–10 deg in diameter and was presented within a circular aperture surrounded by a gray field of mean luminance. Each stimulus orientation was repeated 100 times for each eye. Periods of stimulus presentation lasted 1.28 seconds and were separated by 1.5 s intervals during which we presented a homogeneous gray screen of mean luminance. In one of the amblyopic animals (4 separate array implants) and the control animal, we presented the drifting sinusoidal gratings at 12 orientations and 3 contrast levels (100%, 50%, 12%). In these cases, stimuli were presented for 1 second and each stimulus orientation was repeated 50 times at each of three contrasts. The spatial frequency (1.3 c/deg) and drift rate (6.25 Hz) values for the grating stimuli were chosen to correspond to the typical preference of parafoveal V1 neurons (*DeValois et al. 1982; Foster et al. 1985; Smith et al. 2002*) and to be well within the spatial frequency range where we could behaviorally demonstrate contrast sensitivity in both eyes.

*Spike sorting and analysis criteria.* Our spike sorting procedures have been described in detail previously (Smith and Kohn 2008). In brief, waveform segments exceeding a threshold (based on a multiple of the r.m.s. noise on each channel) were digitized at 30 kHz and stored for offline analysis. We first employed an automated algorithm to cluster similarly shaped waveforms (*Shoham et al. 2003*) and then manually refined the algorithm’s output for each electrode. This manual process took into account the waveform shape, principal component analysis, and inter-spike interval distribution using custom spike sorting software written in Matlab (https://github.com/smithlabvision/spikesort). After offline sorting, we computed a signal to noise ratio metric for each candidate unit (*Kelly et al. 2007*) and discarded any candidate units with SNR below 2.75 as multi-unit recordings. We kept all neurons for which the best grating stimulus evoked a response of more than 2 spikes/second for either the fellow or amblyopic eye. We considered the remaining candidate waveforms (240 units total across sessions) to be high-quality, well isolated single units and we included these units in all further analyses.

### Fano factor

The Fano factor (FF) is defined as across-trial spike count variance divided by mean spike count. We calculated the mean and variance of spike counts for each neuron across 50 repeat trials of an identical high contrast stimulus (stimuli of each orientation were considered as a separate group of 50 repeats). For each neuron-stimulus group of 50 trials, we calculated the mean and variance of spike counts in 100-ms time windows starting at stimulus onset (time 0) and sliding every 50 ms until 850 ms post-stimulus onset. For example, for a time bin of 0-100 ms relative to stimulus onset, counts were made within that 100-ms window at the beginning of each of the 50 trials of each neuron-stimulus pairing, and the mean and the variance were calculated from the resulting set of 50 numbers.

Measurements of the Fano factor are known to be influenced by variability in firing rates: the Fano factor declines as the mean firing rate increases. It is important to take this into account when comparing Fano factor at different time points throughout the trial or for different behavioral conditions to ensure that any significant differences in FF are not simply a consequence of large changes in mean firing rate (*Churchland et al. 2010*). To control for the possible effect of changing firing rates on FF measurements, we used a “mean-matching” method which keeps the population distribution of mean firing rates (but not variances) constant across the analyzed time points and eye stimulation conditions (see *Churchland et al. 2010*). For each eye condition, the mean-matching algorithm first assembled a scatter of the mean rate for each neuron-stimulus set of trials plotted against the variance for each neuron-stimulus pairing, doing so at each time bin. Then, the algorithm selected the greatest common distribution of mean rates across the time points and eye conditions. Then, independently at each time point, neuron-stimulus data points were randomly eliminated if they fell outside the common distribution, and thus not considered in FF calculation for that time point for each eye condition. Importantly, for each eye condition, different neuron-stimulus data points were eliminated, but an equal number of data points remained in subdistributions for the two eye conditions after the elimination. FF was then computed for each eye condition from the remaining neuron-stimulus points as the slope of the regression relating the variance to the mean. The elimination procedure was repeated 10 times, and the resulting FF value for each time point and eye condition was an average of the 10 iterations. We adapted the code provided in the “Variance Toolbox” for MATLAB by M.M. Churchland to do the mean-matching procedure across behavioral conditions in addition to across time points.

### Measures of correlation

Here we provide a brief description of correlation analyses performed for this study. A detailed discussion can be found in two previous publications (*Kohn and Smith 2005; Smith and Kohn 2008*). The r_sc_, also known as spike count correlation or noise correlation, captures the degree to which trial-to-trial fluctuations in responses are shared by two neurons. Quantifying the magnitude of the correlation in trial-to- trial response variability is achieved by computing the Pearson correlation coefficient of evoked spike counts of two cells to many presentations of an identical stimulus. For each session, we paired each neuron with all of the other simultaneously recorded neurons, but excluded any pairs of neurons from the same electrode. We then combined all the pairs from all of the recording sessions in the amblyopic animals, and separately, the control animal. This resulted in 4630 pairs across the 3 amblyopic animals and 155 pairs in one control animal. For each stimulus orientation, we normalized the response to a mean of zero and unit variance (Z-score), and calculated r_sc_ after combining response z-scores across all stimuli. We removed trials on which the response of either neuron was > 3 SDs different from its mean (*Zohary et al. 1994*) to avoid contamination by outlier responses. We also compared our measures of response correlation to the tuning similarity of the two neurons, which we calculated as the Pearson correlation between the mean response of each cell to each of the tested orientations (termed r_signal_). For neurons with similar orientation tuning r_signal_ is closer to 1, while neurons with dissimilar tuning have r_signal_ values approaching −1.

### Curve fitting

We fit the raw data in Figure 4C with the equation:

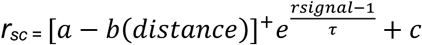

in order to estimate the parameters a (y-intercept), b (slope), τ (exponential decay constant) and c (baseline value). We used the Matlab function *lsqcurvefit,* with initialization parameters based on the fit parameters estimated for r_sc_, r_signal_ and distance data in our previous work in visually normal animals (*Smith and Kohn 2008*). The utilized initialization values were: a = 0.225, b = 0.048, T = 1.87, c = 0.09.

**Figure 4.**
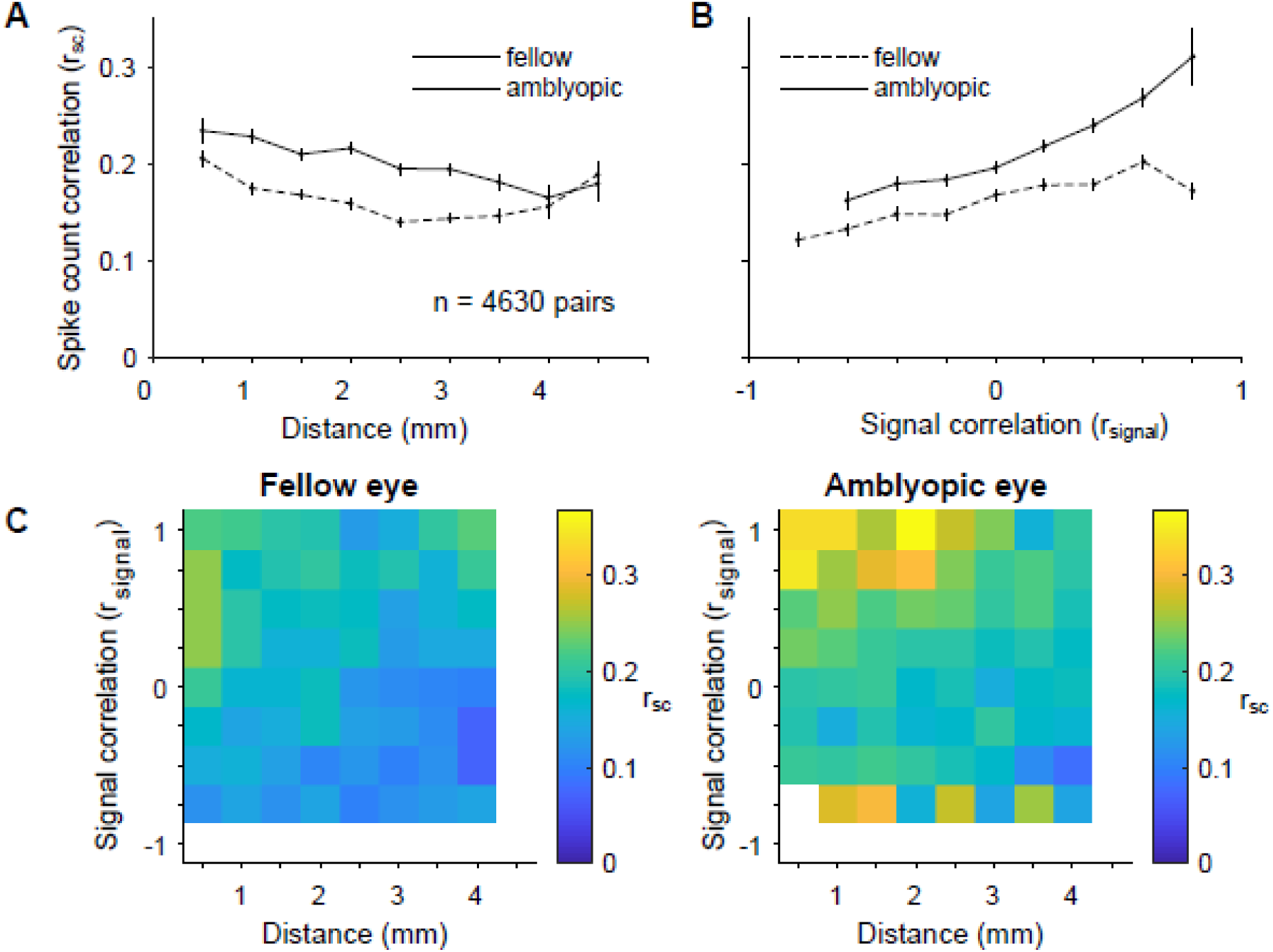
Dependence of rsc on distance and tuning similarity in amblyopic V1. (A) Stimuli presented to the amblyopic eye (solid line) resulted in higher spike count correlation over all possible distances between recorded neurons, as compared to fellow eye stimulation (dashed line). Mean spike count correlation is plotted as a function of the distance between the array electrodes that contain the neurons in each assessed pair. The distance bins start at 0 mm and extend to 4.5 mm in 0.5 mm increments. The average of the rsc values for neuronal pairs included in each bin is plotted at the end value for each bin. Error bars represent ± 1 SEM. (B) For fellow and amblyopic eye stimulation, mean spike count correlation is plotted as a function of signal correlation, which can be thought of as similarity in orientation tuning of the two neurons. The rsignal bins start at −1.0 and extend to 1.0 in 0.2 increments. The average of the rsc values for neuronal pairs included in each bin is plotted at the start value for each bin. As has been reported previously, spike count correlation increased with signal correlation. Furthermore, for the amblyopic eye, the relationship between rsc and rsignal was significantly stronger compared to the fellow eye (p<0.05), indicating that similarly tuned neurons exhibit the largest increase in shared trial-to-trial variability. Error bars represent ± 1 SEM (C) Summary color maps illustrate the relationships between distance, spike count correlation and signal correlation for fellow vs. amblyopic eye stimulation. The scale of the colors is indicated by the bar on the right. rsignal bins start at −1 and extend to 1 in 0.25 increments.

### Ocular dominance analysis

For each unit, we first obtained the average firing rate response to each of the 12 orientations of high contrast gratings, then subtracted the baseline firing rate measured during the interstimulus intervals. Next, we determined each unit’s eye preference by comparing the maximum mean response elicited by visual stimulation of the fellow eye (R_f_) with the same unit’s maximum response to visual stimulation of the amblyopic eye (R_a_). Specifically, we computed an ocular dominance index (ODI) defined as ODI = (R_f_ – R_a_)/(R_f_ + R_a_). The ODI values ranged from −1 to 1, with more negative values signifying a cell’s preference for amblyopic eye stimulation, and more positive values indicating a preference for the fellow eye. For the pairwise analyses, we measured the difference between the ODI values of the cells constituting each pair, such that cells with a very similar eye preference had an ODI difference close to 0, and cells preferring opposite eyes had an ODI difference close to 2.

### Statistical significance tests

All indications of variation in the graphs and text are standard errors of the mean (SEM), unless otherwise noted. The statistical significance of results was evaluated with paired t-tests, unless otherwise noted.

We used a bootstrapping method for statistical testing of the relationships between r_sc_ and r_signal_. Specifically, for 1000 iterations, we sampled with replacement from a pool of matched r_sc_ and r_signal_ values computed for each pair of neurons, separately for each eye condition. Using the “polyfit” function in Matlab, we then computed the slope of a line fit through the scatter of r_sc_ values plotted against the corresponding r_signal_ values for the neuronal pairs used on each sampling iteration. Thus, for each eye stimulation condition, we collected 1000 estimates of the slope of the linear relationship between r_sc_ and r_signal_. We then looked at confidence interval bounds to test for a statistically significant difference between the bootstrapped distributions of slope values computed for amblyopic vs. fellow eye stimulation. We also performed the same bootstrapping procedure to assess whether the relationship between r_sc_ and eye preference was significantly different between fellow and amblyopic eye conditions. We used non-smoothed data for this statistical analysis.

We also used bootstrapping for statistical testing of the inter-ocular difference in delta r_sc._ Briefly, we calculated Δr_sc_ in our data set by subtracting the high contrast r_sc_ value of each neuronal pair from the low contrast r_sc_ value attained for the same pair of neurons. We then performed 1000 iterations of randomly sampling with replacement from the pool of pairs of neurons (1381 pairs total). Each pair of neurons was associated with a high contrast and low contrast r_sc_ value that we could use to compute Δr_sc._ For each eye condition, on each iteration, we computed the average of the sample of Δr_sc_ values. In the end we collected a distribution of 1000 average Δr_sc_ values for each eye condition. We compared these distributions of Δr_sc_ values using confidence interval bounds.

### Decoding stimulus orientation

Within 4 separate recording sessions, we randomly subdivided the spiking data in our two eye conditions such that a subset of the trials was used to train the classifier and the held-out trials were used to assess classification performance. We did 3 rounds of cross-validation such that 3 different random subsets of trials were used for training the classifier. For 3 of the recording sessions (JS 579 and EM 640), we show the average classification performance of 20 classifiers each trained and tested on the responses of 30 randomly selected V1 neurons in each session. In the fourth session (subject HN 580), we only recorded from 30 neurons in total, and thus for this session we assessed performance of just one classifier from 3 rounds of cross-validation. For each round of cross-validation that we performed for each group of 30 neurons, we calculated the classification accuracy of the trained classifier as the proportion of held-out, testing trials that were correctly classified - meaning these trials were assigned their true class labels by the classifier. The remaining three of the total seven sessions had comparatively few simultaneously recorded cells (∼10) and thus were not included in this decoding analysis.

As we had a total of 12 stimulus orientations, for each testing trial, a trained multi-class classifier was tasked with deciding which one of 12 orientations (classes) was most fitting given the V1 population activity on that trial. We used the Error-Correcting Output Coding method (ECOC) which decomposed our multi-class classification problem into many binary classification tasks solved by binary SVM classifiers. In the ECOC framework, the final decision about the class label for a piece of data is achieved by considering the output or “vote” of each subservient binary classifier.

## Results

The overall goal of our study was to examine whether neuronal interactions are altered within primary visual cortex of strabismic amblyopes. To this end, we recorded from populations of V1 neurons using 100-electrode “Utah” arrays while a visual stimulus was separately presented to the amblyopic or the fellow, non-amblyopic eye of anesthetized macaque monkeys. We then evaluated the strength and pattern of correlation in the recorded populations in order to determine if functional interactions among neurons differed during visual stimulation of each eye.

### Behavioral deficits in amblyopic monkeys

Prior to the neural recordings, we characterized the behavioral extent of the amblyopic visual deficits by constructing spatial contrast sensitivity functions for each eye in the amblyopic animals. The fitted curves were used to estimate the optimal spatial frequency and peak contrast sensitivity. For the three strabismic amblyopes, reduced contrast sensitivity and spatial resolution in the amblyopic eye was evident from the reduced peak and spatial extent of the fitted curve (Fig 1). The control animal was tested binocularly and confirmed to be visually normal (Fig 1). Based on these behavioral assessments, we concluded that all three of our experimental animals had severe strabismic amblyopia.

**Figure 1.**
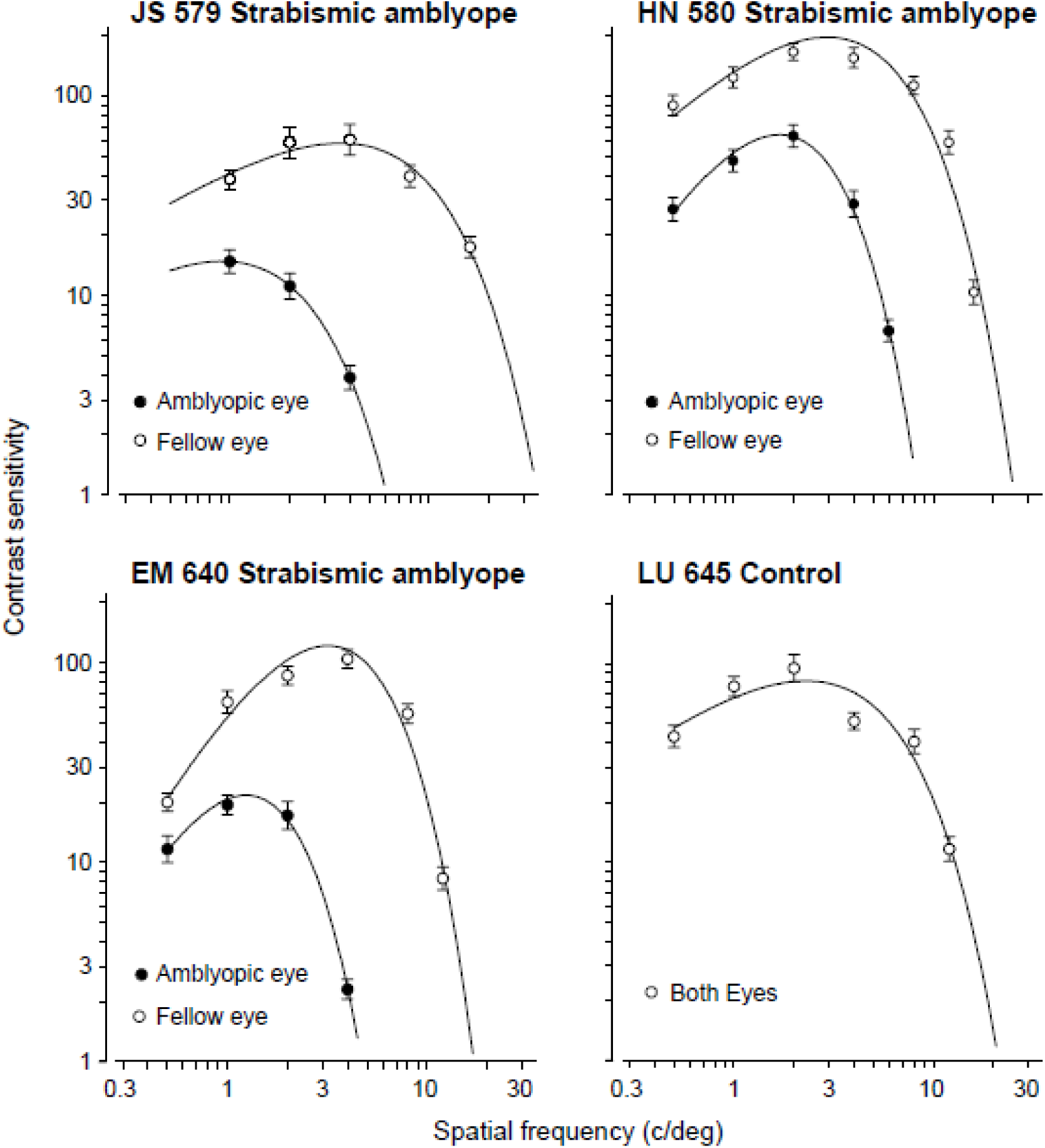
Spatial contrast sensitivity functions, plotted separately for the amblyopic eye (filled symbols) and fellow eye (unoperated, normal eye; open symbols). The four panels show plots for 3 strabismic amblyopes and 1 control, visually normal animal. Behavioral sensitivity loss in the amblyopic eye was observed for all 3 amblyopes: the peak contrast sensitivity was both decreased and shifted to lower spatial frequencies for the amblyopic eyes compared to the fellow eyes.

### Amblyopia affects individual neuronal responsivity

We first studied the changes in single neuron responses in amblyopic primary visual cortex. We recorded from “Utah” arrays while a drifting sinusoidal grating was presented to either the fellow or amblyopic eye of an anesthesized monkey. We presented full-contrast gratings of 12 different orientations to either the amblyopic or fellow eye of three monkeys. For comparison, we also analyzed neural responses to the full-contrast stimuli shown to the right or left eye of the control animal.

We found that most V1 neuronal firing rates were substantially lower during amblyopic eye stimulation compared to fellow eye stimulation (Fig 2A-B). Over the whole population of recorded neurons, the mean maximum spike count across 1-second stimuli presented to the fellow eye was 15.08 ± 1.1 sp/s, compared to 9.56 ± 0.96 sp/s for the same 1-second stimuli presented to the amblyopic eye (p<0.0001, Fig 2B). In the control animal, considering all the recorded neurons, there was no statistically significant difference in maximum evoked firing rates for left versus right eye stimulation (Fig 2C, 9.61 ± 1.67 vs. 9.65 ± 1.55 sp/s, p=0.92).

**Figure 2.**
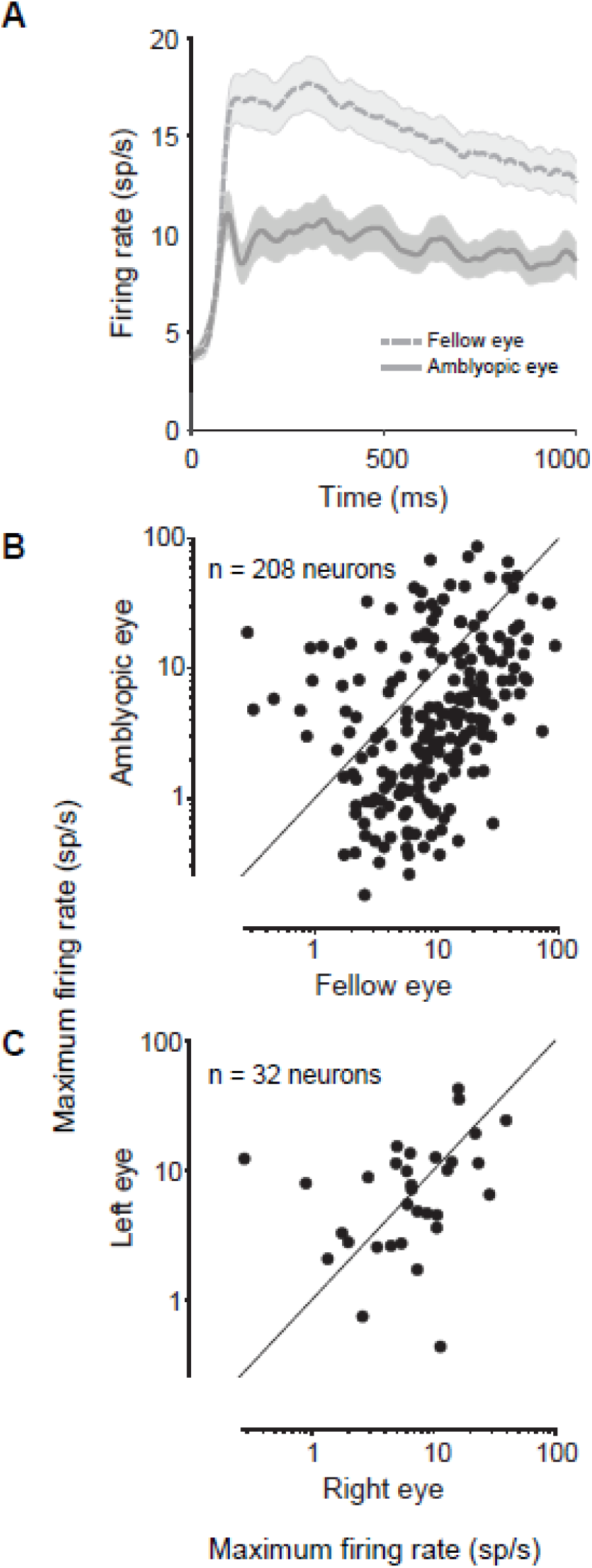
Comparison of neuronal firing rates in response to normal and amblyopic eye stimulation. (A) Peristimulus time histograms (PSTHs) show the population average responses to fellow (dashed line) and amblyopic (solid line) eye stimulation. For each neuron, we computed a PSTH for the one stimulus orientation that evoked the highest response from that neuron, then we averaged across all recorded neurons. Shading represents ± 1 SEM (n = 208 neurons). Neuronal firing rates were greatly diminished upon amblyopic eye stimulation. (B) Each point in the scatter diagram represents the maximum firing rate (spike count during 1 second of stimulus presentation) of each recorded neuron across 12 tested orientations of drifting gratings. The maximum firing rates in response to stimulation of the fellow eye are plotted against the maximum firing rates evoked by amblyopic eye stimulation. The majority of recorded neurons showed decreased responsivity to amblyopic eye stimulation as compared to fellow eye stimulation. Combined across animals, a total of 208 neurons were recorded from V1 of amblyopic animals. (C) Same as in (B), except data for the control animal are shown. A total of 32 neurons were recorded in the control, visually normal animal. There was no observed difference in the maximum firing rates elicited by stimulation of normal right and left eyes.

### Amblyopia alters both response variability and coordinated population activity in V1

It is well known that both the spontaneous and evoked responses of individual neurons are variable even across repeated trials of identical visual stimulation conditions (*Arieli et al. 1996*; *Tolhurst et al. 1983; Shadlen and Newsome 1998).* Recent neurophysiological studies have found that in many primate visual areas, the ongoing response variability declines with the onset of a stimulus (*Churchland et al. 2010*), suggesting that sensory inputs stabilize cortical activity which could in turn improve the reliability of transmitted sensory information. In amblyopia, it is possible that abnormally increased neuronal response variability during stimulus processing contributes to vision problems (*Levi et al. 2008*). In fact, a recent study compared the amount of spiking noise between V2 neurons of amblyopic and visually normal animals, and found that response variability was increased in amblyopic V2 during spontaneous activity and for low contrast visual stimulation (*Wang et al. 2017*).

We quantified whether trial-to-trial response variability of individual neurons in V1 differs between amblyopic and fellow eyes by measuring the Fano factor (FF), or the variance-to-mean ratio, for spiking responses elicited by high contrast stimulation of each eye. Importantly, we utilized a mean matching procedure in our calculation of FF, where we used different subgroups of neurons across different time points and eye stimulation conditions to keep the mean firing rates constant (see Methods). This method ensured that the computed FF values were independent of the any large changes in firing rates between the eye stimulation conditions, or over the course of stimulus presentation.

We assessed the temporal evolution of FF throughout the stimulus duration by calculating FF in 100 ms time windows at multiple time points over the 1 second stimulus. We found that for both eye conditions, there was a sharp decrease in FF after stimulus onset that was consistent with the previously observed time course of FF in a study of numerous cortical areas (Fig 3A; *Churchland et al. 2010*). However, we observed that FF for amblyopic eye stimulation remained significantly higher than FF for fellow eye throughout the whole stimulus duration (Fig 3A), indicating that a high level of spiking variability persists in V1 neurons during processing of visual stimuli presented to the amblyopic eye.

**Figure 3.**
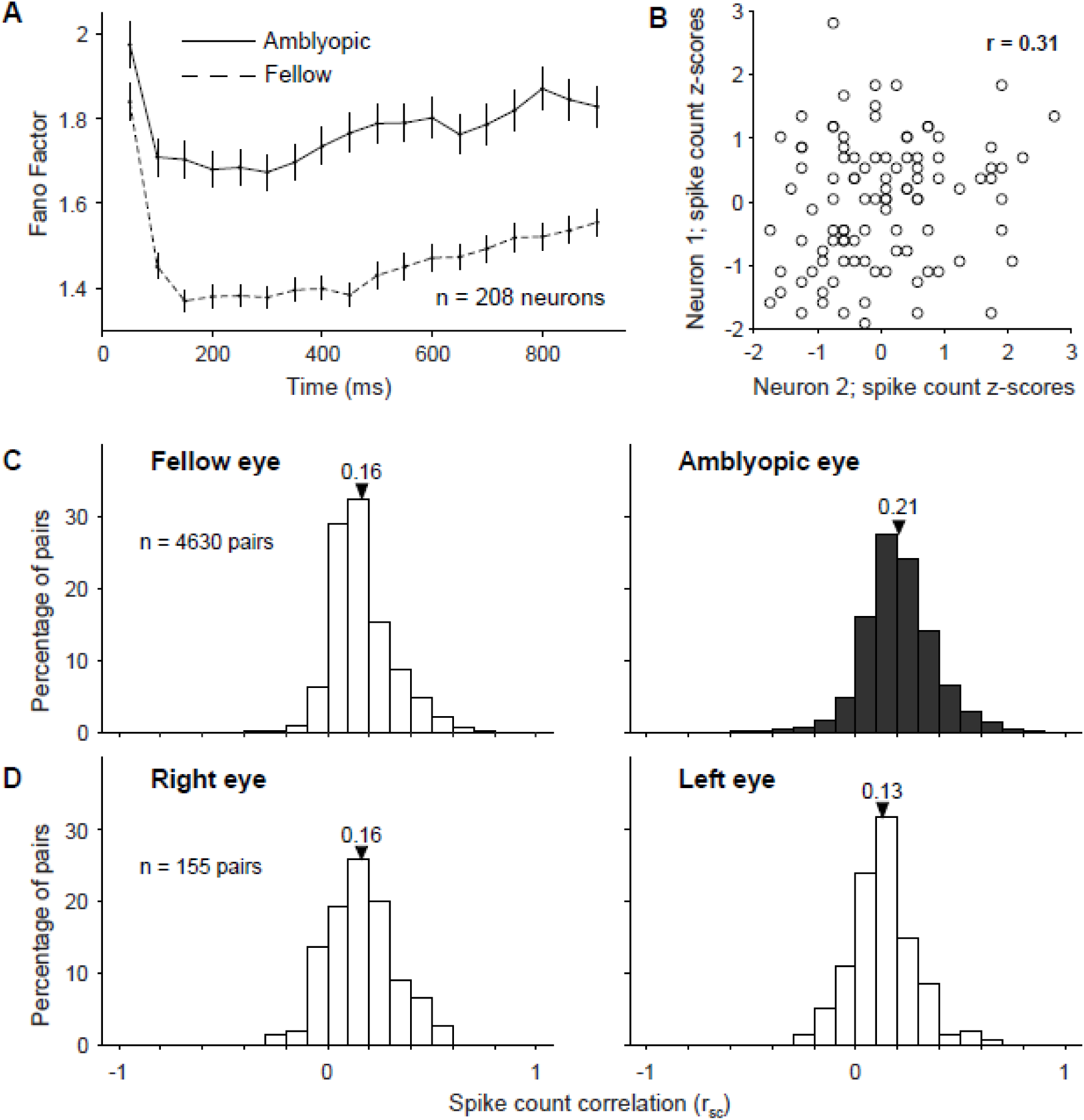
Effect of amblyopia on individual and shared variability of responses to full contrast stimuli in a population of V1 neurons. (A) Mean-matched Fano factor is increased for amblyopic compared to fellow eye stimulation at different time points throughout stimulus presentation. Error bars represent 95% confidence intervals. (B) The scatter plot shows the aggregate, z-transformed, single trial responses of an example pair of recorded V1 neurons to 100 repeat presentations of a single identical full contrast stimulus. Both of the neurons’ responses were ‘noisy’, varying from trial to trial. Spike count correlation (rsc), also known as noise correlation, is computed as the Pearson’s correlation coefficient (r) of the responses of two cells to repeated presentations of an identical stimulus. (C) Shown are the distributions of rsc computed across 4630 pairs of neurons. The mean of each rsc distribution is indicated with a triangle. Spike count correlation was computed separately for neuronal responses evoked by visual stimulation of the amblyopic (filled) and fellow (white) eyes. For each neuronal pair, we calculated the rsc after combining response z-scores across all stimulus orientations. Spike count correlation was significantly increased for pairs of neurons responding to amblyopic eye stimulation, compared to fellow eye stimulation (p<0.00001). (D) Same as in (C), except rsc was computed for 155 pairs of neurons in the control, visually normal animal when either the right or the left eye was stimulated. We did not observe a statistically significant difference in average rsc between right and left eyes of the control animal (p = 0.06).

A small portion of the individual neuron response variability, or noise, is known to be shared between neighboring neurons in cortex. Numerous recent studies have been devoted to understanding how stimulus information is embedded in the population code. In particular, the pattern of correlated variability and its dependence on the stimulus-response structure have been shown in theoretical studies to have potential importance for the information in the population code (*Averbeck et al. 2006; Kohn et al. 2016*). We reasoned that amblyopia could alter the activity pattern and level of interaction in networks of V1 neurons, and might thereby influence information encoding and behavioral performance.

We measured the correlated variability of neural responses to quantify the interactions in pairs of simultaneously recorded V1 neurons. The degree to which trial-to-trial fluctuations in responses are shared by two neurons can be quantified by computing the Pearson correlation of spike count responses to many presentations of the same stimulus (termed spike count correlation, r_sc_, or noise correlation). In Figure 3B, the scatter plot depicts z-transformed spike count responses of two example recorded V1 neurons to an identical stimulus presented to the fellow eye on many trials. The depicted pair of neurons has a positive r_sc_ of 0.31, indicating that responses of these two neurons tend to fluctuate up and down together across trials. We measured correlations over the entire stimulus window (1 second), for all pairs of neurons recorded either during amblyopic or fellow eye stimulation (*see Methods)*.

Correlations for pairs of neurons were significantly larger when a stimulus was presented to the amblyopic eye compared to the fellow eye (Fig 3C; mean r_sc_ 0.21 (0.17 SD) vs mean r_sc_ 0.16 (0.14 SD); p<0.00001). Because we randomized the visual stimulus between the eyes across trials, we were able to make this comparison directly in the same neurons. This difference in r_sc_ between amblyopic and fellow eye stimulation provides evidence for altered functional interactions in the same population of neurons. Furthermore, our finding of a higher (mean matched) Fano factor for amblyopic compared to fellow eye stimulation suggests that the changes in covariance among the V1 neuron responses must be quite large, leading to increased noise correlations despite a concomitant increased variance of individual neuronal responses to amblyopic eye stimulation. There was no apparent difference in r_sc_ between the stimulation of the right and the left eyes in the control animal (Fig 3D; mean r_sc_ 0.16 (0.17 SD) vs mean r_sc_ 0.13 (0.15 SD); p=0.06). In both the control and amblyopic animals, our recordings were targeted to the superficial layers of V1, where previous studies have reported r_sc_ values higher than in the intermediate and deep layers (*Hansen et al. 2013; Smith et al. 2013*). The distribution of r_sc_ values we observed across our animals is consistent with the range of values reported by previous studies using similar and different recording preparations in V1 of primates (*Gutnisky and Dragoi 2008; Kohn and Smith 2005; Reich et al. 2001; Smith and Kohn 2008; see Cohen and Kohn 2011* for an extensive summary of previously observed r_sc_ values).

### Stimulus-dependent correlation structure is modified in amblyopic V1

Several experimental and theoretical studies suggest that the structure of correlations – the dependence of correlations on the functional properties and physical location of neurons – can have a strong influence on the information encoded by the population (*see Averbeck et al. 2006; Kohn et al. 2016 for reviews*). Previous work in normal macaque V1 and V4 has shown that correlations are highest for pairs of neurons that are near each other and that have similar orientation tuning preferences *(Kohn and Smith 2005; Ruff and Cohen 2016; Smith and Kohn 2008; Smith and Sommer 2013).* Here, we investigated whether the correlation structure observed in visual cortex of normal animals is maintained in the cortex of amblyopes. To do this, we first examined if r_sc_ measurements differed depending on the distance between the neurons in each pair. We found that r_sc_ was largest for pairs of neurons near each other, compared to pairs of neurons farther apart, for both fellow and amblyopic eye stimulation (Fig 4A & C). Thus, for cortical processing of visual information received through the amblyopic eye, correlations were increased for all pairs of neurons, regardless of the distance between them.

We next investigated whether the relationship between tuning similarity and the magnitude of correlations was altered in the cortex of amblyopes. We used sinusoidal gratings of 12 different orientations to engage neurons with varied orientation preferences, which enabled us to assess the tuning similarity of each pair of neurons. Tuning similarity was quantified by calculating r_signal_, the Pearson correlation of the mean responses of two neurons to each of 12 stimulus orientations. To test how functional interactions varied among neurons with different tuning preferences, we calculated r_sc_ as a function of r_signal_. As in previous studies, we found that r_sc_ was highest for neurons with similar tuning (positive r_signal_), and lowest for neurons with opposite tuning preferences (negative r_signal_), for both fellow and amblyopic eye stimulation (Fig 4B). However, for the amblyopic eye, the relationship between r_sc_ and r_signal_ was significantly stronger compared to the fellow eye (p < 0.05; *see Methods* for details of bootstrapping and statistical testing), such that pairs of similarly tuned neurons exhibited the largest difference in r_sc_ between the amblyopic and fellow eye stimulation conditions (Fig 4B&C). That is, pairs of similarly tuned neurons show the largest increase in r_sc_ between fellow and amblyopic eye stimulation. So, both raw correlation for stimulation of each eye as well as the difference in correlation between activity evoked by stimulation of the two eyes depend on tuning similarity of a pair of neurons. In the control animal, we found that r_sc_ was highest for neurons with similar tuning and lowest for neurons with opposite tuning preferences, for both left and right eye stimulation, as previously reported in normal animals.

The summary color maps in Figure 4C depict the dependence of r_sc_ on distance and r_signal_ for amblyopic and fellow eye visual stimulation. In a previous study of V1 neurons in visually normal animals, we found that the dependence of r_sc_ on both cortical distance and tuning similarity is well characterized by a product of two functions:

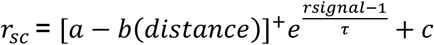

where the linear term represents the decay of r_sc_ with distance, the exponential decay represents how r_sc_ declines with r_signal_, and []^+^ indicates that negative values of the linear terms are set to 0 (*Smith and Kohn 2008*). We fit the data from our amblyopic animals in Figure 4C with this same equation. For fellow eye condition, the linear decay had an intercept (a) of 0.121 ± 0.038 (95% confidence interval), and a slope (b) of 0.048 ± 0.02 mm^-1^ while the exponential decay constant (*τ*) was 0.936 ± 0.47 (unitless) and the baseline (c) added to the product of the functions was 0.149 ± 0.006. For the amblyopic eye, the linear decay had an intercept (a) of 0.267 ± 0.055 (95% confidence interval), and a slope (b) of 0.038 ± 0.019 mm^-1^ while the exponential decay constant (*τ*) was 0.67 ± 0.3 and the baseline (c) added to the product of the functions was 0.151 ± 0.026. The intercept, slope and baseline values for both of the eye conditions were similar to those reported for V1 neurons of normal animals in our previous work (*Smith and Kohn 2008*). This similarity indicates that the relationship between distance and r_sc_ in amblyopic animals of this study is not altered compared to normal animals of our previous study. On the other hand, the value of *τ* was lower for the amblyopic animals of this study compared to the value (1.87 ± 0.67) reported in our previous work in normal animals. A smaller value of *τ* indicates that the rate at which r_sc_ values decline as r_signal_ values decrease is faster in amblyopes, which is consistent with our analysis of the relationship between r_sc_ and r_signal_ in Figure 4B. Overall, our results suggest that amblyopia affects not only the overall level of correlation, but also the extent to which neurons interact with their neighbors of both similar and dissimilar stimulus preferences.

### Increased correlations predominate among amblyopic V1 neurons that preferentially respond to fellow eye

In strabismic amblyopic monkeys, binocular organization in V1 is disrupted, such that the ocular dominance distribution becomes U-shaped with a significant reduction in binocularly activated cells (*Baker et al. 1974; Kiorpes et al. 1998; Smith et al. 1997; Wiesel 1982)*. Additionally, several studies report a decrease in the number of cortical neurons that preferentially respond to visual stimulation through the amblyopic over the fellow eye (*Hubel and Wiesel 1965; Crawford and von Noorden 1979; Kiorpes et al. 1998; Movshon et al. 1987; Shooner et al. 2015; (in cat) Schröder et al. 2002)*. Specific changes in the circuitry underlying the eye preference and binocular responsivity of V1 neurons could be reflected in an altered pattern of pairwise interactions in the population. Therefore, we next examined whether our observed changes in spike count correlation were associated with eye preference changes of individual neurons in amblyopic V1.

For each cell, we first computed an ocular dominance index (ODI) as a measure of the cell’s eye preference. ODI distributions in each amblyopic animal ranged between the values of −1 and 1, with more negative and positive values indicating higher responsivity to visual stimuli viewed through the amblyopic or fellow eye, respectively. Figure 5A shows a distribution of ODI values for 208 neurons recorded from the 3 amblyopic animals. We observed an ocular dominance bias toward positive values, indicating that the majority of cells fired more strongly in response to visual stimulation of the fellow eye than the amblyopic eye (141 neurons with ODI value > 0.2 and 36 neurons with ODI value < −0.2). There were relatively few binocularly activated V1 neurons in our amblyopic animals (31 neurons with ODI values within +/- 0.2 of 0).

**Figure 5.**
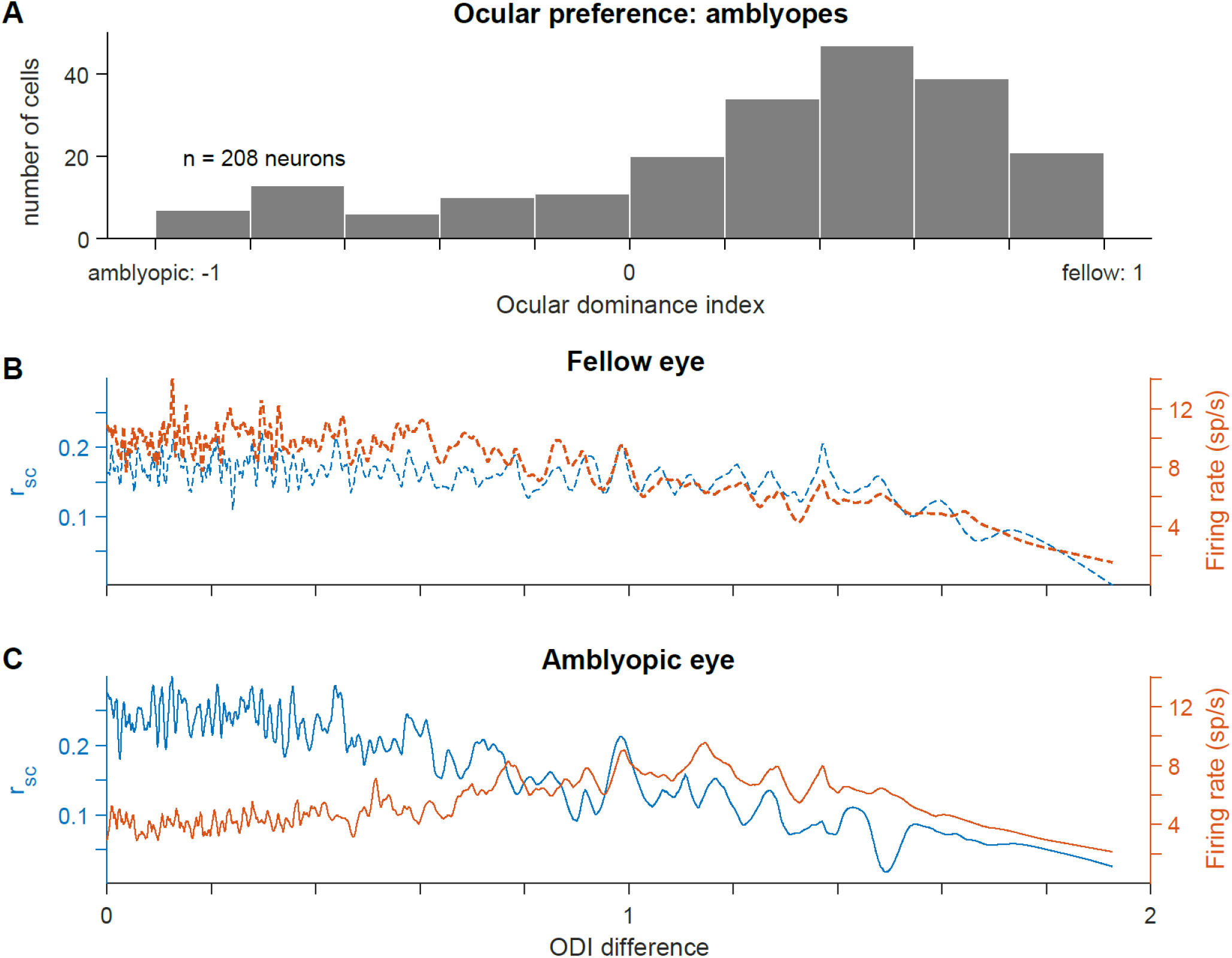
Relationship between ocular dominance changes and increased correlations in amblyopic V1. (A) A histogram showing the ocular dominance index (ODI) values for all 208 neurons recorded across the 3 amblyopic animals. Neurons with ODI values closer to −1 preferentially responded to visual input through the amblyopic eye, while neurons with ODI values closer to 1 had higher respons ivity to fellow eye visual stimulation. The ODI values were unevenly distributed, and biased toward the fellow eye (ODI < −0.2: 36 neurons; - 0.2<ODI<0.2: 31 neurons; ODI>0.2: 141 neurons). (B) For fellow eye visual stimulation, spike count correlation values (left y-axis) and firing rates (right y-axis) are plotted as a function of the difference in ODI values of the neurons in each pair. An ODI difference closer to 0 indicates that the neurons composing the pair have the same ocular preference. The traces shown were produced by smoothing over the data points with a sliding window (size of window = 15 data points). (C) same as in (B), but considering V1 responses to visual stimulation through the amblyopic eye. Neurons with similar ODIs had higher correlations during amblyopic eye stimulation, compared to the level of correlations in the same neuron pairs during fellow eye stimulation (p < 0.05).

We next investigated whether the magnitude of spiking correlations was dependent on the eye from which each neuron received its dominant input. In this analysis, we measured correlations in pairs of neurons as a function of the difference in eye preference between the cells in each pair, termed ODI difference. Differences in ODI ranged from 0 to 2, where cells that preferred the same eye had an ODI difference of 0, while cells that preferred opposite eyes had an ODI difference of 2. Because of the ocular dominance bias in our neuronal population, the majority of neuronal pairs with an ODI difference close to 0 preferred the fellow eye. We first analyzed the magnitude of correlation as a function of the ODI difference, and found that there was a negative relationship in both the fellow (Fig 5B) and amblyopic (Fig 5C) eye, indicating that pairs of neurons that preferred the same eye had higher correlations than pairs of neurons that had opposite eye preferences. This effect could be due simply to the lower mean firing rates among pairs of neurons that preferred quite dissimilar stimuli. For the fellow eye, this was indeed the case – the correlation tracked the geometric mean firing rate of the pairs of neurons. However, for the amblyopic eye there was a particularly high level of correlation among neurons that preferred input from the same eye (ODI difference < 0.8) that could not be explained by the firing rates. When comparing the same pairs of neurons under different eye stimulation conditions, the neuronal pairs with an ODI difference < 0.8 had decreased responsivity but higher correlations during amblyopic eye stimulation, compared to fellow eye stimulation. Accordingly, we found that the relationship between eye preference similarity and the magnitude of correlations in pairs of neurons was significantly different between the two eyes (stronger for the amblyopic eye, p<0.05; *see Methods* for details on bootstrapping and statistical testing). These results indicate that in amblyopia there is not only a weaker representation of the amblyopic eye at the single neuron level in V1, as has been shown before, but also that the ocular dominance changes in individual neurons are related to changes in functional interactions among those neurons.

### Decoding stimulus orientation from amblyopic V1 population activity

The modifications in pattern and strength of functional interactions that we observed in amblyopic V1 could degrade the encoding of stimuli presented to the amblyopic eye. Therefore, we compared how well the recorded network of V1 neurons represented stimulus information when high contrast visual input was delivered through the amblyopic versus the fellow eye. We used a statistical classification method to decode stimulus orientation from the activity of simultaneously recorded V1 neurons (see *Methods* for details). As we had a total of 12 stimulus orientations, for each testing trial, a trained multi-class classifier was tasked with deciding which one of 12 possible classes was most consistent with the V1 population activity on that trial. Using this classification analysis, we explored whether visual stimulus information was harder to read out from V1 population activity when the amblyopic eye provided the input.

We found that classification accuracy was substantially decreased when a classifier was trained and tested on neuronal responses during amblyopic eye stimulation compared to training and testing on V1 responses to fellow eye stimulation. Figure 6 shows decoding accuracy for fellow versus amblyopic eye stimulation trials for four different recording sessions across 3 animals. While decoding performance remained above chance (8.33%) for both of the eyes in all four examined sessions, accuracy was consistently reduced when decoding from neural responses to amblyopic eye visual input. Importantly, classification performance is dependent on the response properties and orientation tuning of the recorded neuronal population. For instance, we observed different decoding accuracies for two recording sessions that were conducted in the same animal (JS579) because the sampling of neurons was different.

**Figure 6.**
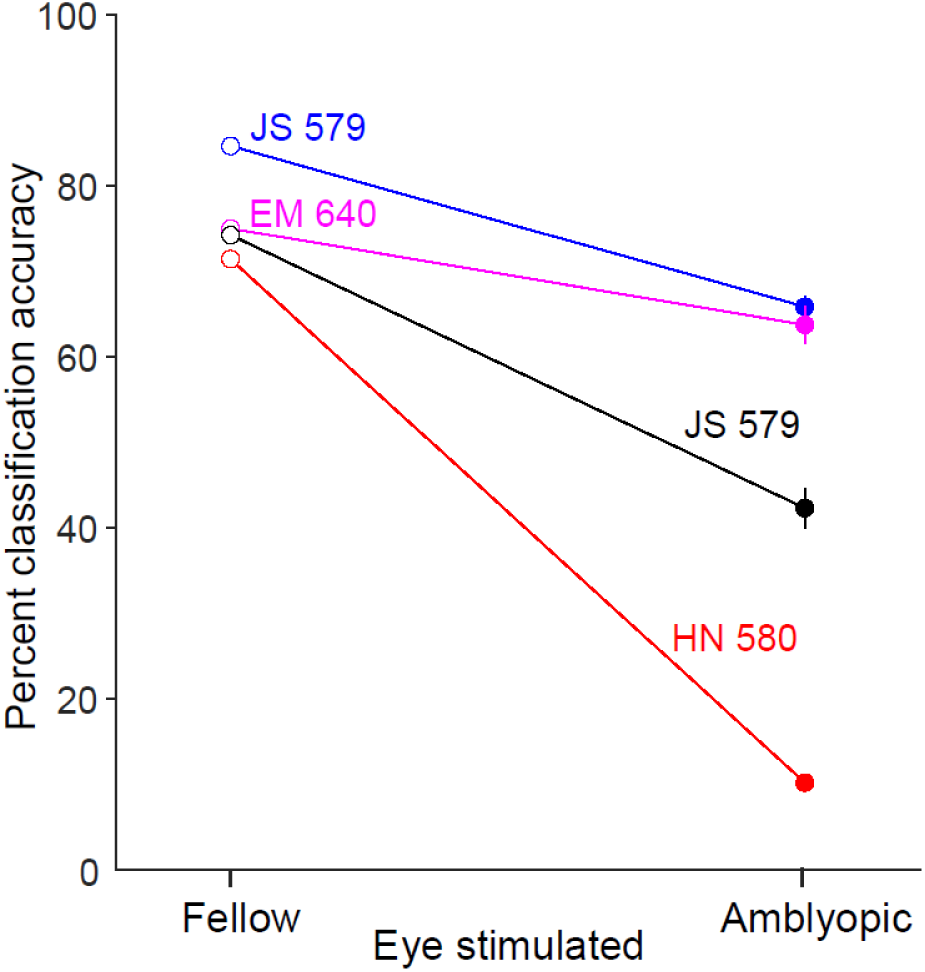
Decoding grating orientation from fellow or amblyopic eye stimulation. When trained and tested on neuronal responses during amblyopic eye stimulation, the decoding accuracy was decreased compared to when a decoder is trained and tested on responses to fellow eye stimulation. The four colors correspond to decoding results from neuronal responses on 4 different array implants across 3 animals.

### Effect of stimulus contrast on correlated variability in amblyopic V1

Despite previous work, our understanding of the neural basis for diminished contrast sensitivity in amblyopes remains incomplete. It is possible that in amblyopia, a deficit in global network responsivity to contrast is more pronounced than individual neuron response deficits. Importantly, studies in visually normal animals have shown that stimulus contrast can affect the level of interactions in a neuronal population. For instance, correlations in pairs of V1 neurons depend on stimulus contrast, such that r_sc_ is significantly larger for low contrast stimuli than high contrast stimuli (*Kohn and Smith 2005*). This suggests that spontaneous cortical activity has a considerable amount of inherent correlated variability which can be reduced by strong stimulus drive *(Churchland et al. 2010; Smith and Kohn 2008; Snyder et al. 2014)* Developmental abnormalities in the visual cortex of amblyopes could affect how networks of cortical neurons interpret the strength of stimulus drive provided by high vs. low contrast stimuli. Based on these observations in normal animals, we wondered how the amount of stimulus drive to the amblyopic eye affects the strength of correlated variability in V1.

We presented full (100%), medium (50%) and low (12%) contrast gratings of 12 different orientations, separately to the amblyopic or fellow eye of one of the amblyopic monkeys. We then measured the correlation in response variability of 1381 neuronal pairs in the recorded neuronal population for each stimulus contrast presented to each of the two eyes. Because r_sc_ values for neuronal pairs are known to depend on the firing rates of constituent neurons (see *Cohen and Kohn 2011*), for this analysis, we binned the computed r_sc_ values by geometric mean firing rate of neuronal pairs. This method allowed us to study the effect of stimulus contrast on correlated variability in amblyopic V1 while accounting for the wide range of responsivity observed across the recorded individual neurons (Fig 2B).

In agreement with the results of *Kohn and Smith (2005),* when we analyzed the V1 population response on trials with fellow eye stimulation, lowering stimulus contrast significantly increased mean r_sc_ for all neural pairs regardless of their geometric mean firing rate (Fig 7A). Interestingly, for stimuli presented to the amblyopic eye, r_sc_ was relatively insensitive to the level of contrast (Fig 7B). That is, a full contrast stimulus viewed by the amblyopic eye did not substantially reduce the amount of correlated variability in most V1 neurons (except those with very high firing rates) compared to a lower contrast stimulus. This is apparent when viewing a contrast response function for correlation (Fig 8), where the relatively flat lines in low-firing rate pairs of neurons for amblyopic eye stimulation indicate a lack of contrast sensitivity of correlation.

**Figure 7.**
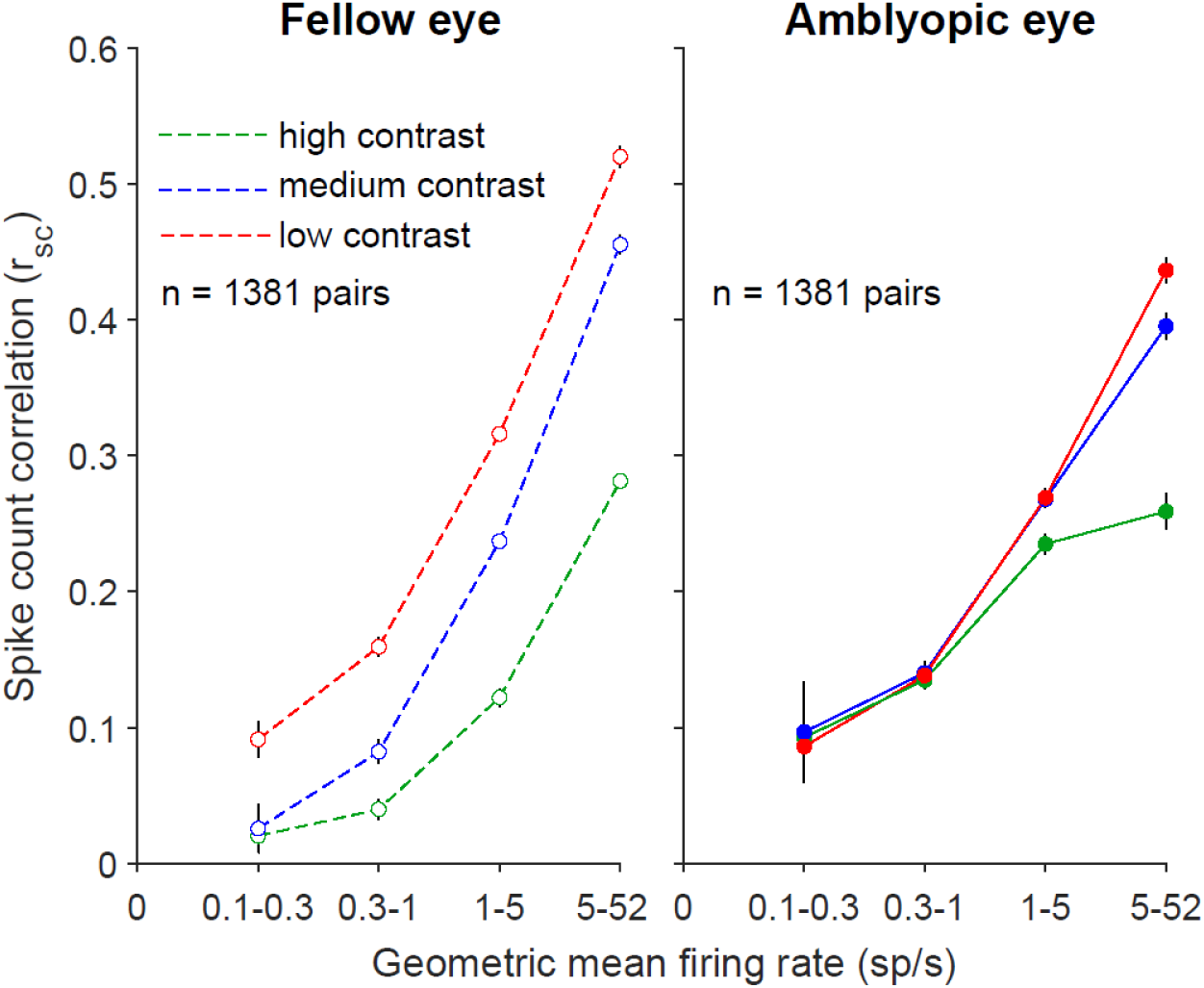
The average of the rsc values for neuronal pairs in each geometric mean firing rate bin is plotted, for grating stimuli of high (green, 100%), medium (blue, 50%), and low (red, 12%) contrasts. Error bars represent s.e.m. For the fellow eye, lowering stimulus contrast significantly increased mean rsc at all firing rates, while with amblyopic eye stimulation, rsc was relatively unaffected by stimulus contrast. Computing the difference in rsc between high and low contrast (Δrsc) for all 1381 neuron pairs revealed a significant inter-ocular disparity in Δrsc in the amblyopic animal (p<0.05; based on confidence intervals of bootstrapped, mean Δrsc distributions).

**Figure 8.**
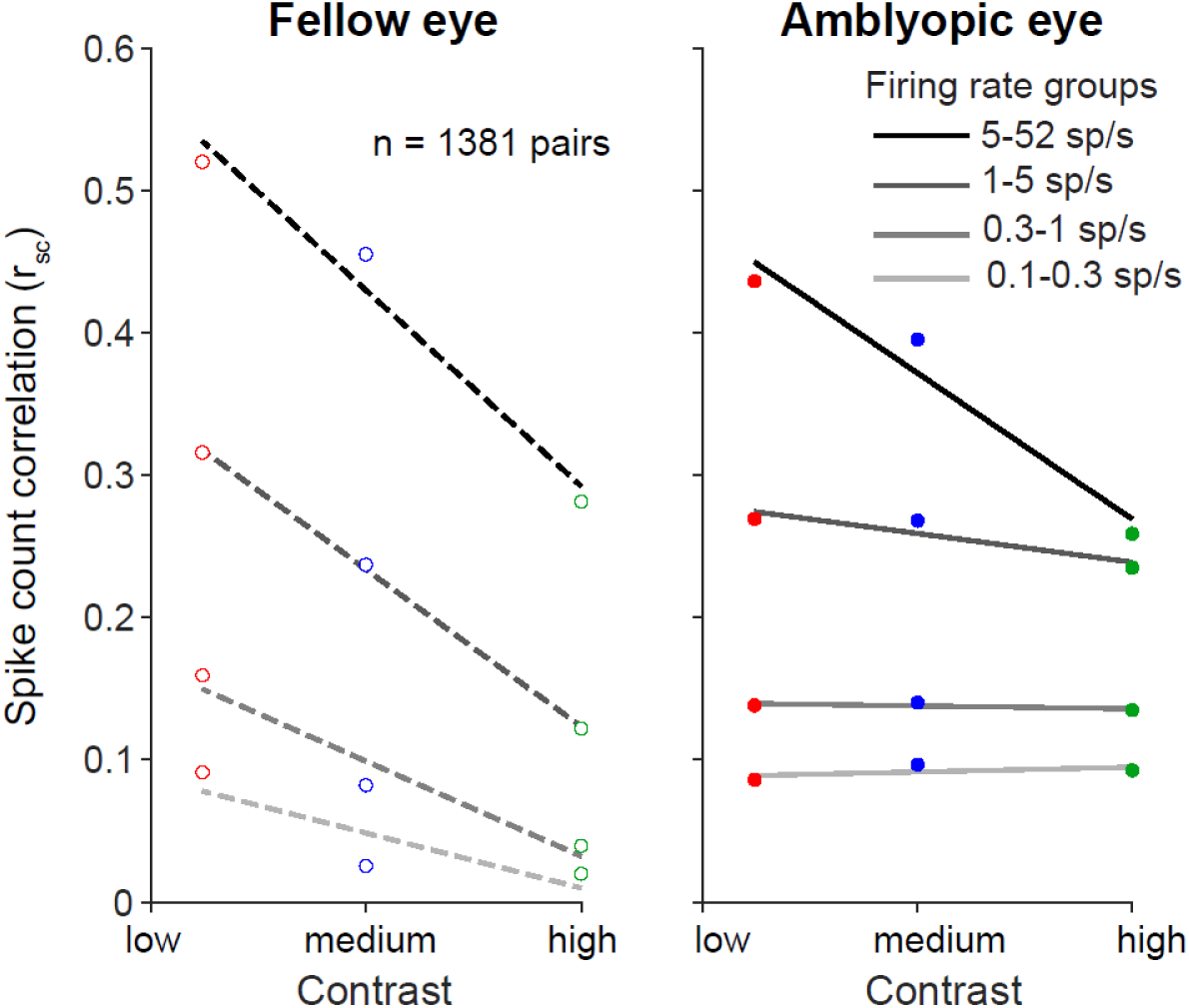
Dependence of spike count correlation on stimulus contrast. Amblyopic eye stimulation resulted in similar rsc across three stimulus contrasts (100%, 50% and 12%). rsc values are binned according to the mean firing rate for each neuronal pair, and the average rsc value per firing rate bin is plotted as a function of contrast.

We observed that the mean r_sc_ values for high firing neuronal pairs responding to fellow eye stimulation were higher than the r_sc_ values in the highest firing rate bins for the amblyopic eye condition (Fig 7). Because some neurons in our population retained high firing rates to stimuli shown to the amblyopic eye, it is expected that the r_sc_ values for neuronal pairs in the high firing rate bin would be more similar to those for fellow eye. Additionally, although most neurons we recorded had a significantly higher r_sc_ for amblyopic than fellow eye stimulation, the ocular preferences of the neurons can play a role in how responsive the neurons are to each eye, and thus can influence the relative difference in r_sc_ magnitude between amblyopic and fellow eyes. For instance, if there are two neurons that have a slight preference for the right (fellow) eye, they will have higher firing rates (and a higher r_sc_ value) in response to right (fellow) eye visual stimulation compared to left (amblyopic) eye visual stimulation. In such a scenario, the effect of increased r_sc_ during amblyopic eye stimulation would not be as apparent.

We next quantified the differential effect of stimulus contrast on the amount of correlated variability for the fellow versus the amblyopic eye. For each neuron pair, we computed the difference in r_sc_ between high and low contrast (Δr_sc_) for each eye condition. Since Δr_sc_ is computed by subtracting high contrast r_sc_ values from low contrast r_sc_ values, the closer Δr_sc_ is to 0, the more similar are the r_sc_ values computed during high and low contrast stimulation. This metric revealed that indeed, the Δr_sc_ distribution for amblyopic eye stimulation was shifted closer to 0, and was significantly different from the Δr_sc_ distribution computed for fellow eye stimulation (amblyopic mean = −0.1017, fellow mean = −0.1523; p<0.05; based on confidence intervals of bootstrapped, mean Δr_sc_ distributions). Furthermore, we also found a significant difference in the strength of this interocular disparity between the amblyopes and the control animal (p<0.0001). Thus, for stimulus processing through the amblyopic eye, neurons had not only impaired contrast sensitivity measured one cell at a time (*Kiorpes et al. 1998; Movshon et al. 1987*), but also maintained high levels of correlated variability even in the presence of strong stimulus input.

## Discussion

Our goal in this study was to gain insight into the neural basis of amblyopia by looking for abnormalities beyond those already known to affect individual neuronal responses. We recorded simultaneously from tens of neurons in the primary visual cortex of monkeys with strabismic amblyopia, which allowed us to measure the functional interactions between pairs of neurons during visual stimulation of the fellow, non-amblyopic eye versus the amblyopic eye of each animal. Our primary finding was that the structure of correlated trial-to-trial response variability among V1 neurons was altered in amblyopic compared to fellow eye stimulation. Specifically, stimulation of the amblyopic eye resulted in stronger correlations that were restricted to neurons with similar orientation tuning and similar eye preference, and these correlations were relatively insensitive to stimulus drive. To examine the consequence of these changes for stimulus representation in networks of amblyopic V1 neurons, we decoded grating orientation from simultaneously recorded populations of neurons. The accuracy of decoding stimulus orientation for amblyopic eye stimulation was reduced compared to decoding the same stimuli from neural activity in response to fellow eye stimulation. Taken together, these results demonstrate profound shifts in the functional response properties and interactions among neurons in amblyopic cortex when the stimulus is presented to the amblyopic eye.

### Altered circuitry in V1 of amblyopes

What do our observed differences in r_sc_ between the two eyes suggest about circuits of V1 neurons that process visual information received from amblyopic eye? To answer this question, it is first necessary to consider the physiological sources of correlated variability (*for review see Doiron et al. 2016*). Correlations in pairs of neurons are thought to arise in part from common afferent projections to the two neurons (*Shadlen and Newsome 1998*). Correlations can also arise from feedback (top down) signals (*Cumming and Nienborg 2016*), feedforward processing of stimuli (*Kanitscheider et al. 2015*), recurrent connectivity in local circuits (*Doiron et al. 2016*), and from variable synaptic transmission due to the dynamics of vesicle release *(Doiron et al. 2016*). Changes in correlated variability may therefore reflect reorganization in the underlying circuitry, and correlation analysis has previously proved useful for assessing changes in functional connectivity (*Cohen and Newsome 2008; Greschner et al. 2011; Reid and Alonso 1995*).

In our study of amblyopic V1, we found that during amblyopic eye stimulation, there was elevated pairwise correlation in V1 neuronal responses, and that this remained unchanged across low, medium and high stimulus drive to the amblyopic eye. Our results suggest that in amblyopic visual systems, networks of V1 neurons have altered connectivity and function abnormally when processing visual information received through the amblyopic eye. In particular, our observation that increased correlation persists across a range of stimulus intensities shown to the amblyopic eye suggests that V1 neurons may not fully engage in processing stimulus information received through an amblyopic eye. Previous studies measuring individual neuronal contrast response functions have found that few amblyopic V1 neurons have reduced contrast sensitivity at high spatial frequencies, and that the observed reduction in neuronal contrast sensitivity is not enough to account for contrast perception deficits found in amblyopic animals (*Kiorpes et al. 1998; Movshon et al. 1987; see also: Shooner et al. 2015*). However, a recent study (*Wang et al. 2017*) found that contrast response functions for V2 neurons responding to amblyopic eye stimulation in anisometropic amblyopes were abnormal. Our findings indicate that amblyopia-related contrast processing deficits could manifest both downstream of V1 and at the level of neuronal correlations in V1.

According to our results, it is likely that visual stimuli received through the amblyopic eye have a weaker influence in the visual cortex due to both single-neuron and network level changes following a shift in ocular dominance towards the fellow eye. In the amblyopic animals of this study, the majority of the recorded V1 neurons preferentially responded to stimulus drive through the fellow eye, and there were few binocularly responsive neurons. Furthermore, the difference in correlated variability and firing rates between amblyopic and fellow eye stimulation was restricted to pairs of cells that had the same eye preference. Together, these results are consistent with a re-wiring scheme in which a substantial portion of the neurons lose amblyopic eye inputs but gain or retain fellow eye inputs during abnormal visual experience. Anatomically, the representation of the amblyopic eye in pairs of V1 neurons could decline as a result of altered lateral connections in V1, from reduced thalamocortical projections that carry amblyopic eye information, or both. Studies of horizontal connections in amblyopic macaques and cats have reported reduced connectivity between cells located in opposite ocular dominance columns in the superficial layers of V1, but connectivity between neurons in columns dominated by the same eye is normal (*Löwel and Singer 1992 (cat); Löwel 1994 (cat); Trachtenberg and Stryker 2001 (cat); Tychsen et al. 1992, 1997, 2004 (macaque)*). At a coarse level, the structure of thalamocortical inputs remains largely normal in amblyopic monkeys (*Adams et al. 2015; Fenstemaker et al. 2001; Hendrickson et al. 1987; Horton et al. 1996*). But even with structurally intact thalamocortical projections, the effectiveness of thalamocortical drive to V1 could be reduced specifically for inputs from the amblyopic eye if there were changes in how cortical circuits receive and process these inputs. To that point, we recently described local circuit changes in V1, in particular, reduction in excitatory drive to amblyopic eye neurons resulting in a change in E/I balance, that could explain the abnormal response to contrast variation during amblyopic eye viewing (*Hallum et al. 2017; Shooner et al. 2015, 2017*).

When considering changes across the entire population of neurons, it is evident that the effect of amblyopia is heterogenous across the V1 population. For instance, although most neurons exhibited a higher level of correlations and lower firing rates for amblyopic eye stimulation, a subgroup of neurons retained normal responsivity and continued to respond well to stimulation of the amblyopic eye. Specifically, neuronal pairs with the highest firing rates did not show an increase in correlation compared to the same high firing neuronal pairs responding to fellow eye stimulation (Figs 5 and 7). This observation is consistent with prior reports that some neurons in amblyopic cortex retain normal response properties. For example, some neurons in amblyopic cortex in monkeys maintained high responsivity to high spatial frequencies while other neurons had altered responsivity (*Kiorpes et al. 1998; Movshon et al. 1987*). This co-existence of normally responsive and altered cells in amblyopic V1 highlights the importance of considering pairwise interactions in the context of the properties of the cells in each pair, which can reveal subgroups of neurons (and types of visual stimulus information) that are particularly affected.

### Decoding information from V1 populations

A number of studies suggest that correlated variability between sensory neurons might be especially important for encoding of stimulus information in populations of neurons (*Abbott and Dayan 1999; Averbeck et al. 2006; Cohen and Maunsell 2009; Cohen and Kohn 2011*). Furthermore, there is some evidence for a direct link between changes in correlated variability and shifts in psychophysical performance (*Beaman and Dragoi 2017; Cohen and Maunsell 2009; Zohary et al. 1994*). Importantly, not only the amount of correlated variability in a given network, but also the particular neurons that have altered interactions, matters for stimulus representation. Here, we found that the increase in correlations was highest for pairs of similarly tuned neurons. A common finding of theoretical and experimental studies is that an increase in amount of shared noise between similarly tuned neurons is detrimental for population coding (*Averbeck et al. 2006; Ecker et al. 2011; Jeanne et al. 2013).* Our results thus indicate that stimulus representation is degraded in populations of V1 neurons that process visual stimuli shown to the amblyopic eye, and that this effect is greater than would be expected simply from the reduced responses observed in individual neurons.

Our decoding analysis demonstrates that, as expected, stimulus information is harder to read out from V1 population activity when amblyopic eye rather than the fellow, non-amblyopic eye provides the visual input. Classification accuracy was consistently reduced when decoding stimulus orientation from neural responses to amblyopic compared to fellow eye stimulation. This is consistent with the idea that stimulus representation in V1 is impaired for amblyopic eye signals, which can in turn lead to downstream errors in information processing. Interestingly, amblyopic observers have global perceptual deficits that are not simply predicted by single neuron changes in V1 (*Kozma and Kiorpes 2003*). For instance, strabismic amblyopes have impaired performance in contour integration, a task that requires identifying a curve imbedded in a noisy background (*Kozma and Kiorpes 2003; Levi et al. 2007)*. In this study we found a larger increase in correlations between similarly tuned neurons compared to neurons with dissimilar tuning during amblyopic eye stimulation. Perhaps deficits in contour integration in amblyopia arise from decreased accuracy in coordinating V1 representations of neighboring, similarly oriented pieces of the contour. Overall, our findings indicate that to more conclusively define the neurophysiological correlates of visual deficits in amblyopia, it is important to consider population-level processing of visual information and not just the properties of single neurons.

### Theories for the neural basis of amblyopia

Previous work provides evidence for at least four neurophysiological correlates of amblyopic visual deficits, including 1) altered responsivity and tuning of single neurons in V1, 2) neural changes in visual areas downstream of V1, 3) reduced cortical representation of the amblyopic eye (“undersampling”) and 4) topographical jitter, or disorder in neural map of visual space (*Kiorpes et al. 1998; Kiorpes 2006, 2016; Levi 2013; Wang et al. 2017).* In this study we found that the strength and pattern of functional interactions in pairs of neurons in the primary visual cortex was different when processing amblyopic eye and fellow eye inputs. We therefore conclude that abnormalities in visual representation at the level of V1 neuron populations may constitute a fifth factor contributing to amblyopic visual deficits. Further work will be needed to determine the relative contributions of these factors to amblyopic visual losses.

## Acknowledgements

KA was supported by a National Science Foundation (NSF) Graduate Fellowship Grant 1747452, MAS was supported by National Institutes of Health (NIH) grants R00EY018894, R01EY022928, R01MH118929, R01EB026953, P30EY008098, NSF grant NCS 1734901, a career development grant and an unrestricted award from Research to Prevent Blindness, and the Eye and Ear Foundation of Pittsburgh. LK, JAM, and the creation and testing of the amblyopic subjects were supported by NIH grant R01EY05864 to LK and P51 OD010425 to the Washington National Primate Research Center. We are grateful to Michael Gorman for his assistance rearing and behaviorally testing animals, to Howard M. Eggers for creating experimental strabismus, and to Romesh Kumbhani, Najib Majaj, Yasmine El-Shamayleh and others in the Movshon laboratory for their assistance during recording experiments.

## References

1. Abbott LF, Dayan P. The effect of correlated variability on the accuracy of a population code. Neural Comput 11(1):91–101, 1999.

2. Adams DL, Economides JR, Horton JC. Contrasting effects of strabismic amblyopia on metabolic activity in superficial and deep layers of striate cortex. J Neurophysiol 113(9):3337–44, 2015.

3. Arieli A, Sterkin A, Grinvald A, Aertsen A. Dynamics of ongoing activity: explanation of the large variability in evoked cortical responses. Science 273(5283):1868–71, 1996.

4. Asper L, Crewther D, Crewther SG. Strabismic amblyopia: Part1. Psychophysics. Clin Exp Optom 83(4):200–11, 2000.

5. Averbeck BB, Latham PE, Pouget A. Neural correlations, population coding and computation. Nat Rev Neurosci 7(5):358–66, 2006.

6. Baker DH, Meese TS, Hess RF. Contrast masking in strabismic amblyopia: attenuation, interocular suppression and binocular summation. Vision Res 48(15):1625–40, 2008.

7. Baker FH, Grigg P, von Noorden GK. Effects of visual deprivation and strabismus on the response of neurons in the visual cortex of the monkey, including studies on the striate and prestriate cortex in the normal animal. Brain Research 66(2):185–208, 1974.

8. Beaman CB, Eagleman SL, Dragoi V. Sensory coding accuracy and perceptual performance are improved during the desynchronized cortical state. Nat Commun 8(1):1308, 2017.

9. Bi H, Zhang B, Tao X, Harwerth RS, Smith EL, Chino YM. Neuronal responses in visual area V2 (V2) of macaque monkeys with strabismic amblyopia. Cereb Cortex 21(9):2033–45, 2011.

10. Blakemore C, Vital-Durand F. Effects of visual deprivation on the development of the monkey’s lateral geniculate nucleus. J Physiol (Lond) 380:493–511, 1986.

11. Bradley A, Freeman RD. Contrast sensitivity in anisometropic amblyopia. Invest Ophthalmol Vis Sci 21(3):467–76, 1981.

12. Chino YM, Shansky MS, Jankowski WL, Banser FA. Effects of rearing kittens with convergent strabismus on development of receptive-field properties in striate cortex neurons. J Neurophysiol 50(1):265–86, 1983.

13. Churchland MM, Yu BM, Cunningham JP, et al. Stimulus onset quenches neural variability: a widespread cortical phenomenon. Nat Neurosci 13(3):369–78, 2010.

14. Cohen MR, Maunsell JH. Attention improves performance primarily by reducing interneuronal correlations. Nat Neurosci 12(12):1594–600, 2009.

15. Cohen MR, Kohn A. Measuring and interpreting neuronal correlations. Nat Neurosci 14(7):811–9, 2011.

16. Cohen MR, Newsome WT. Context-dependent changes in functional circuitry in visual area MT. Neuron 60(1):162–73, 2008.

17. Crawford ML, de Faber JT, Harwerth RS, Smith EL 3^rd^, von Noorden GK. The effects of reverse monocular deprivation in monkeys. II. Electrophysiological and anatomical studies. Exp Brain Res 74(2):338–47, 1989.

18. Crawford ML, Harwerth RS. Ocular dominance column width and contrast sensitivity in monkeys reared with strabismus or anisometropia. Invest Ophthalmol Vis Sci. 45(9):3036–42, 2004.

19. Crawford ML, Von noorden GK. Concomitant strabismus and cortical eye dominance in young rhesus monkeys. Trans Ophthalmol Soc U K 99(3):369–74, 1979.

20. Crewther DP, Crewther SG. Neural site of strabismic amblyopia in cats: spatial frequency deficit in primary cortical neurons. Exp Brain Res 79(3):615–22, 1990.

21. Cumming BG, Nienborg H. Feedforward and feedback sources of choice probability in neural population responses. Curr Opin Neurobiol 37:126–132, 2016.

22. De valois RL, Yund EW, Hepler N. The orientation and direction selectivity of cells in macaque visual cortex. Vision Res 22(5):531–44, 1982.

23. Doiron B, Litwin-kumar A, Rosenbaum R, Ocker GK, Josić K. The mechanics of state-dependent neural correlations. Nat Neurosci 19(3):383–93, 2016.

24. Ecker AS, Berens P, Tolias AS, Bethge M. The effect of noise correlations in populations of diversely tuned neurons. J Neurosci 31(40):14272–83, 2011.

25. El-Shamayleh Y, Kiorpes L, Kohn A, Movshon JA. Visual motion processing by neurons in area MT of macaque monkeys with experimental amblyopia. J Neurosci 30(36):12198–209, 2010.

26. Farzin F, Norcia AM. Impaired visual decision-making in individuals with amblyopia. J Vis 11(14), 2011.

27. Fenstemaker SB, Kiorpes L, Movshon JA. Effects of experimental strabismus on the architecture of macaque monkey striate cortex. J Comp Neurol 438(3):300–17, 2001.

28. Foster KH, Gaska JP, Nagler M, Pollen DA. Spatial and temporal frequency selectivity of neurones in visual cortical areas V1 and V2 of the macaque monkey. J Physiol (Lond) 365:331–63, 1985.

29. Greschner M, Shlens J, Bakolitsa C, et al. Correlated firing among major ganglion cell types in primate retina. J Physiol (Lond) 589(Pt 1):75–86, 2011.

30. Gu Y, Liu S, Fetsch CR, et al. Perceptual learning reduces interneuronal correlations in macaque visual cortex. Neuron 71(4):750–61, 2011.

31. Gutnisky DA, Dragoi V. Adaptive coding of visual information in neural populations. Nature 452(7184):220–4, 2008.

32. Hallum LE, Shooner C, Kumbhani RD, et al. Altered Balance of Receptive Field Excitation and Suppression in Visual Cortex of Amblyopic Macaque Monkeys. J Neurosci 37(34):8216–8226, 2017.

33. Hamm LM, Black J, Dai S, Thompson B. Global processing in amblyopia: a review. Front Psychol 5:583, 2014.

34. Hansen BJ, Chelaru MI, Dragoi V. Correlated variability in laminar cortical circuits. Neuron. 2012;76(3):590–602.

35. Hendrickson AE, Movshon JA, Eggers HM, Gizzi MS, Boothe RG, Kiorpes L. Effects of early unilateral blur on the macaque’s visual system. II. Anatomical observations. J Neurosci 7(5):1327–39, 1987.

36. Hess RF, Howell ER. The threshold contrast sensitivity function in strabismic amblyopia: evidence for a two type classification. Vision Res 17(9):1049–55, 1977.

37. Horton JC, Hocking DR, Kiorpes L. Pattern of ocular dominance columns and cytochrome oxidase activity in a macaque monkey with naturally occurring anisometropic amblyopia. Vis Neurosci 14(4):681–9, 1997.

38. Hou C, Kim YJ, Lai XJ, Verghese P. Degraded attentional modulation of cortical neural populations in strabismic amblyopia. J Vis 16(3):16, 2016.

39. Hubel DH, Wiesel TN. Binocular interaction in striate cortex of kittens reared with artificial squint. J Neurophysiol 28(6):1041–59, 1965.

40. Jeanne JM, Sharpee TO, Gentner TQ. Associative learning enhances population coding by inverting interneuronal correlation patterns. Neuron 78(2):352–63, 2013.

41. Kanitscheider I, Coen-cagli R, Pouget A. Origin of information-limiting noise correlations. Proc Natl Acad Sci USA 112(50):E6973–82, 2015.

42. Kelly RC, Smith MA, Samonds JM, Kohn A, Bonds AB, Movshon JA, Lee TS. Comparison of recordings from microelectrode arrays and single electrodes in the visual cortex. J Neurosci 27(2):261–4, 2007.

43. Kiorpes L. Visual processing in amblyopia: animal studies. Strabismus 14(1):3–10, 2006.

44. Kiorpes L. The Puzzle of Visual Development: Behavior and Neural Limits. J Neurosci 36(45):11384–11393, 2016.

45. Kiorpes L, Daw N. Cortical correlates of amblyopia. Vis Neurosci 35:E016, 2018.

46. Kiorpes L, Kiper DC, O’keefe LP, Cavanaugh JR, Movshon JA. Neuronal correlates of amblyopia in the visual cortex of macaque monkeys with experimental strabismus and anisometropia. J Neurosci 18(16):6411–24, 1998.

47. Kiorpes L, Tang C, Movshon JA. Factors limiting contrast sensitivity in experimentally amblyopic macaque monkeys. Vision Res 39(25):4152–60, 1999.

48. Kiorpes L, Tang C, Movshon JA. Sensitivity to visual motion in amblyopic macaque monkeys. Vis Neurosci 23(2):247–56, 2006.

49. Kohn A, Coen-cagli R, Kanitscheider I, Pouget A. Correlations and Neuronal Population Information. Annu Rev Neurosci 39:237–56, 2016.

50. Kohn A, Smith MA. Stimulus dependence of neuronal correlation in primary visual cortex of the macaque. J Neurosci 25(14):3661–73, 2005.

51. Kozma P, Kiorpes L. Contour integration in amblyopic monkeys. Vis Neurosci 20(5):577–88, 2003.

52. LeVay S, Wiesel TN, Hubel DH. The development of ocular dominance columns in normal and visually deprived monkeys. J Comp Neurol 191(1):1–51, 1980.

53. Levi DM, Harwerth RS. Spatio-temporal interactions in anisometropic and strabismic amblyopia. Invest Ophthalmol Vis Sci 16(1):90–5, 1977.

54. Levi DM, Yu C, Kuai SG, Rislove E. Global contour processing in amblyopia. Vision Res 47(4):512–24, 2007.

55. Levi DM, Klein SA, Chen I. What limits performance in the amblyopic visual system: seeing signals in noise with an amblyopic brain. J Vis 8(4):1.1–23, 2008.

56. Levi DM. Linking assumptions in amblyopia. Vis Neurosci 30(5-6):277–87, 2013.

57. Löwel S, Singer W. Selection of intrinsic horizontal connections in the visual cortex by correlated neuronal activity. Science 255(5041):209–12, 1992.

58. Löwel S. Ocular dominance column development: strabismus changes the spacing of adjacent columns in cat visual cortex. J Neurosci 14(12):7451–68, 1994.

59. McKee SP, Levi DM, Movshon JA. The pattern of visual deficits in amblyopia. J Vis 3(5):380–405, 2003.

60. Meier K, Giaschi, D. Unilateral amblyopia affects two eyes: fellow eye deficits in amblyopia. Invest Ophthalmol Vis Sci 58:1779–1800, 2017.

61. Meier K, Sum B, Giaschi D. Global motion perception in children with amblyopia as a function of spatial and temporal stimulus parameters. Vision Res 127:18–27, 2016.

62. Mitchell JF, Sundberg KA, Reynolds JH. Spatial attention decorrelates intrinsic activity fluctuations in macaque area V4. Neuron 63(6):879–88, 2009.

63. Movshon JA, Eggers HM, Gizzi MS, Hendrickson AE, Kiorpes L, Boothe RG. Effects of early unilateral blur on the macaque’s visual system. III. Physiological observations. J Neurosci 7(5):1340–51, 1987.

64. Ni AM, Ruff DA, Alberts JJ, Symmonds J, Cohen MR. Learning and attention reveal a general relationship between population activity and behavior. Science 359(6374):463–465, 2018.

65. Pham A, Carrasco M, Kiorpes L. Endogenous attention improves perception in amblyopic macaques. J Vis 18(3):11, 2018.

66. Reich DS, Mechler F, Victor JD. Independent and redundant information in nearby cortical neurons. Science 294(5551):2566–8, 2001.

67. Reid RC, Alonso JM. Specificity of monosynaptic connections from thalamus to visual cortex. Nature 378(6554):281–4, 1995.

68. Rislove EM, Hall EC, Stavros KA, Kiorpes L. Scale-dependent loss of global form perception in strabismic amblyopia. J Vis 10(12):25, 2010.

69. Roelfsema PR, König P, Engel AK, Sireteanu R, Singer W. Reduced synchronization in the visual cortex of cats with strabismic amblyopia. Eur J Neurosci 6(11):1645–55, 1994.

70. Rousche PJ, Normann RA. A method for pneumatically inserting an array of penetrating electrodes into cortical tissue. Ann Biomed Eng 20(4):413–22, 1992.

71. Ruff DA, Cohen MR. Stimulus Dependence of Correlated Variability across Cortical Areas. J Neurosci 36(28):7546–56, 2016.

72. Schröder JH, Fries P, Roelfsema PR, Singer W, Engel AK. Ocular dominance in extrastriate cortex of strabismic amblyopic cats. Vision Res 42(1):29–39, 2002.

73. Shadlen MN, Newsome WT. The variable discharge of cortical neurons: implications for connectivity, computation, and information coding. J Neurosci 18(10):3870–96, 1998.

74. Shoham S, Fellows MR, Normann RA. Robust, automatic spike sorting using mixtures of multivariate t-distributions. J Neurosci Methods 127(2):111–22, 2003.

75. Shooner C, Hallum LE, Kumbhani RD, et al. Population representation of visual information in areas V1 and V2 of amblyopic macaques. Vision Res 114:56–67, 2015.

76. Shooner C, Hallum LE, Kumbhani RD, et.al. Asymmetric dichoptic masking in visual cortex of amblyopic macaque monkeys. J Neurosci 37(36):8734–41, 2017.

77. Smith EL, Chino YM, Ni J, Cheng H, Crawford ML, Harwerth RS. Residual binocular interactions in the striate cortex of monkeys reared with abnormal binocular vision. J Neurophysiol 78(3):1353–62, 1997.

78. Smith MA, Kohn A. Spatial and temporal scales of neuronal correlation in primary visual cortex. J Neurosci 28: 12591–12603, 2008.

79. Smith MA, Bair W, Movshon JA. Signals in macaque striate cortical neurons that support the perception of glass patterns. J Neurosci 22(18):8334–45, 2002.

80. Smith MA, Jia X, Zandvakili A, Kohn A. Laminar dependence of neuronal correlations in visual cortex. J Neurophysiol 109(4):940–7, 2013.

81. Smith MA, Sommer MA. Spatial and temporal scales of neuronal correlation in visual area V4. J Neurosci 33(12):5422–32, 2013.

82. Snyder AC, Morais MJ, Smith MA. Dynamics of excitatory and inhibitory networks are differentially altered by selective attention. J Neurophysiol 116:1807–1820, 2016.

83. Tao X, Zhang B, Shen G, Wensveen J, Smith EL 3rd, Nishimoto S, Ohzawa I, Chino YM. Early monocular defocus disrupts the normal development of receptive field structure in V2 neurons of macaque monkeys. J Neurosci 34(41):13840–54, 2014.

84. Trachtenberg JT, Stryker MP. Rapid anatomical plasticity of horizontal connections in the developing visual cortex. J Neurosci 21(10):3476–82, 2001.

85. Tolhurst DJ, Movshon JA, Dean AF. The statistical reliability of signals in single neurons in cat and monkey visual cortex. Vision Res 23(8):775–85, 1983.

86. Tychsen L, Burkhalter A. 1992. Naturally-strabismic primate lacks intrinsic horizontal connections for binocular vision in striate cortex. Soc Neurosci Abstr 18:1455.

87. Tychsen L, Burkhalter, A. Nasotemporal asymmetries in V1: Ocular dominance columns of infant, adult, and strabismic macaque monkeys. J Comp Neurol 388: 32–46, 1997.

88. Tychsen L, Wong AM, Burkhalter A. Paucity of horizontal connections for binocular vision in V1 of naturally strabismic macaques: Cytochrome oxidase compartment specificity. J Comp Neurol 474(2):261–75, 2004.

89. Wang Y, Zhang B, Tao X, Wensveen JM, Smith EL, Chino YM. Noisy spiking in visual area V2 of amblyopic monkeys. J Neurosci 37:922–935, 2017.

90. Wiesel TN. Postnatal development of the visual cortex and the influence of environment. Nature 299:583–91, 1982.

91. Wiesel TN, Hubel DH. Single-cell responses in striate cortex of kittens deprived of vision in one eye. J Neurophysiol 26:1003–17, 1963.

92. Zhou J, Reynaud A, Yao Z, Liu R, Feng L, Zhou Y, Hess RF. Amblyopic suppression: Passive attenuation, enhanced dichoptic masking by the fellow eye or reduced dichoptic masking by the amblyopic eye? Invest Ophthalmol Vis Sci 59(10):4190–7, 2018.

93. Zohary E, Shadlen MN, Newsome WT. Correlated neuronal discharge rate and its implications for psychophysical performance. Nature 370(6485):140–3, 1994.

